# Genome wide mapping of DNA lesions by Repair Assisted Damage Detection sequencing – RADD-Seq

**DOI:** 10.1101/2021.02.07.430183

**Authors:** Noa Gilat, Dena Fridman, Hila Sharim, Sapir Margalit, Natalie R. Gassman, Yael Michaeli, Yuval Ebenstein

## Abstract

Mapping DNA damage and its repair has immense potential in understanding environmental exposures, their genotoxicity, and their impact on human health. Monitoring changes in genomic stability also aids in the diagnosis of numerous DNA-related diseases, like cancer, and assists in monitoring their progression and prognosis. However, genome-wide maps of DNA damage distribution are challenging to produce. Here we describe the localization of DNA damage and repair loci by Repair Assisted Damage Detection sequencing – RADD-Seq. Based on the enrichment of damage lesions coupled with a pull-down assay and followed by next generation sequencing, this method is easy to perform and can produce compelling results with minimal coverage. RADD-seq enables the localization of both DNA damage and repair sites for a wide range of single-strand damage types. Using this technique, we created a genome-wide map of oxidative DNA damage loci before and after repair. Oxidative lesions were heterogeneously distributed along the human genome, with less damage occurring in tight chromatin regions. Furthermore, we showed repair is prioritized for highly expressed, essential genes and in open chromatin regions. RADD-seq sheds light on cellular repair mechanisms and capable of identifying genomic hotspots prone to mutation.

## Introduction

Cellular DNA is continuously exposed to various exogenous and endogenous damaging agents (Lindahl and Barnes 2000; De Bont and Van Larebeke 2004). Damage accumulation on the DNA backbone or bases has cytotoxic and genotoxic effects, leading to mutations, genomic instability and, consequently, cancer (Friedberg 2003). Hence, DNA repair is vital to the integrity of the genome and the organism’s health and survival. Due to the importance of understanding and monitoring DNA damage and repair processes, various methods have been developed to quantify the overall damage level in a cell population and genomic DNA; these mainly include ELISA, comet assays and immunohistochemistry assays (Wani et al. 1984; Yoshida et al. 2002; Manis et al. 2004; Cosaceanu et al. 2007; Bonner et al. 2008). While these methods allow for determining the extent of damage in a DNA sample in bulk, they fail to provide information regarding the local distribution of genomic DNA damage. Mapping damage along the genome reveals the propensity of different genomic regions to accumulate damage and identifies associations to other genomic features, like transcription sites. While several short-read next-generation sequencing (NGS)-based methods locate and assess damage levels along the genome (Hu et al. 2015; Adar et al. 2016; Hu et al. 2016; Mao et al. 2016; Poetsch et al. 2018; Wu et al. 2018), none presently enables the facile and accurate mapping of DNA damage loci and repair dynamics for various damage types.

Single-strand lesions represent the most common class of damage incurred by DNA molecules, and these lesions are repaired extensively by cellular mechanisms. Specifically, oxidative DNA damage, manifested as single-strand breaks (SSB) and 8-oxo-7,8-dihydroguanine (8-oxoG) modifications, is typically repaired by either the nucleotide excision repair (NER) or the base excision repair (BER) pathways. In both processes, the DNA lesions are removed and replaced in three core steps: (1) the damaged DNA base is removed by a glycosylase, (2) incision of the DNA backbone is performed by an endonuclease or the bifunctional activity of the glycosylase, and (3) repair is performed, by gap filling and ligation of the nicked backbone (Friedberg 2003; Houtgraaf et al. 2006). Here, we introduce a method for mapping single-strand DNA lesions that can identify the genomic regions prone to damage accumulation (“damage hotspots”). The method also detects the loci favorable for repair and reveal their repair dynamics. This method, termed Repair Assisted Damage Detection sequencing (RADD-seq), uses repair enzymes to excise the DNA lesions leaving a single-strand gap. The damage sites are then repaired *in-vitro* with biotinylated nucleotides, followed by DNA fragmentation, immunoprecipitation and sequencing. Reads are mapped back to the reference genome, indicating the locations of damage sites.

To demonstrate the capability of our method to map and monitor repair dynamics, the osteosarcoma cell line U2OS was exposed to the oxidizing agent potassium bromate, KBrO_3_. RADD-seq was then used to monitor DNA lesion induction and repair in a time-dependent manner. For this demonstration, we map the landscape of DNA oxidative damage using the human 8-OxoGuanine Glycosylase 1 (hOGG1) repair enzyme to specifically excise oxidative DNA lesions. However, RADD-seq is compatible with any repair enzyme of interest, allowing any desired damage type to be monitored. We found that KBrO_3_-induced oxidative DNA damage was heterogeneously distributed along the genome. Moreover, we show that DNA repair occurs more readily in genomic regions with higher levels of gene expression and open chromatin than low expression, closed chromatin genomic regions. Finally, we identified specific gene clusters in which repair tends to occur more extensively, presumably due to their role in cell viability.

## Results

RADD-seq exploits the incorporation of biotinylated nucleotides into DNA damage sites to provide a quick and straightforward pull-down based method for genome-wide sequencing DNA damage loci (**Fig. 1**). Using extracted genomic DNA, base lesions are recognized and excised from the double-strand by a specific repair enzyme, leaving single-strand gaps (**Fig. 1A**,**B**). Next, a DNA polymerase fills the gaps using biotinylated nucleotides, incorporating these nucleotides into the original damage sites (**Fig. 1C**). DNA is then sheared into ~150 bp fragments by sonication and immunoprecipitated using anti-biotin antibodies, enriching the sample for damaged DNA fragments (**Fig. 1D**). Illumina sequencing is conducted and reads are mapped back to a reference genome. Genomic coverage is then examined to identify trends along the genome, such as damage hotspots (**Fig. 1E**).

**Figure 1.**
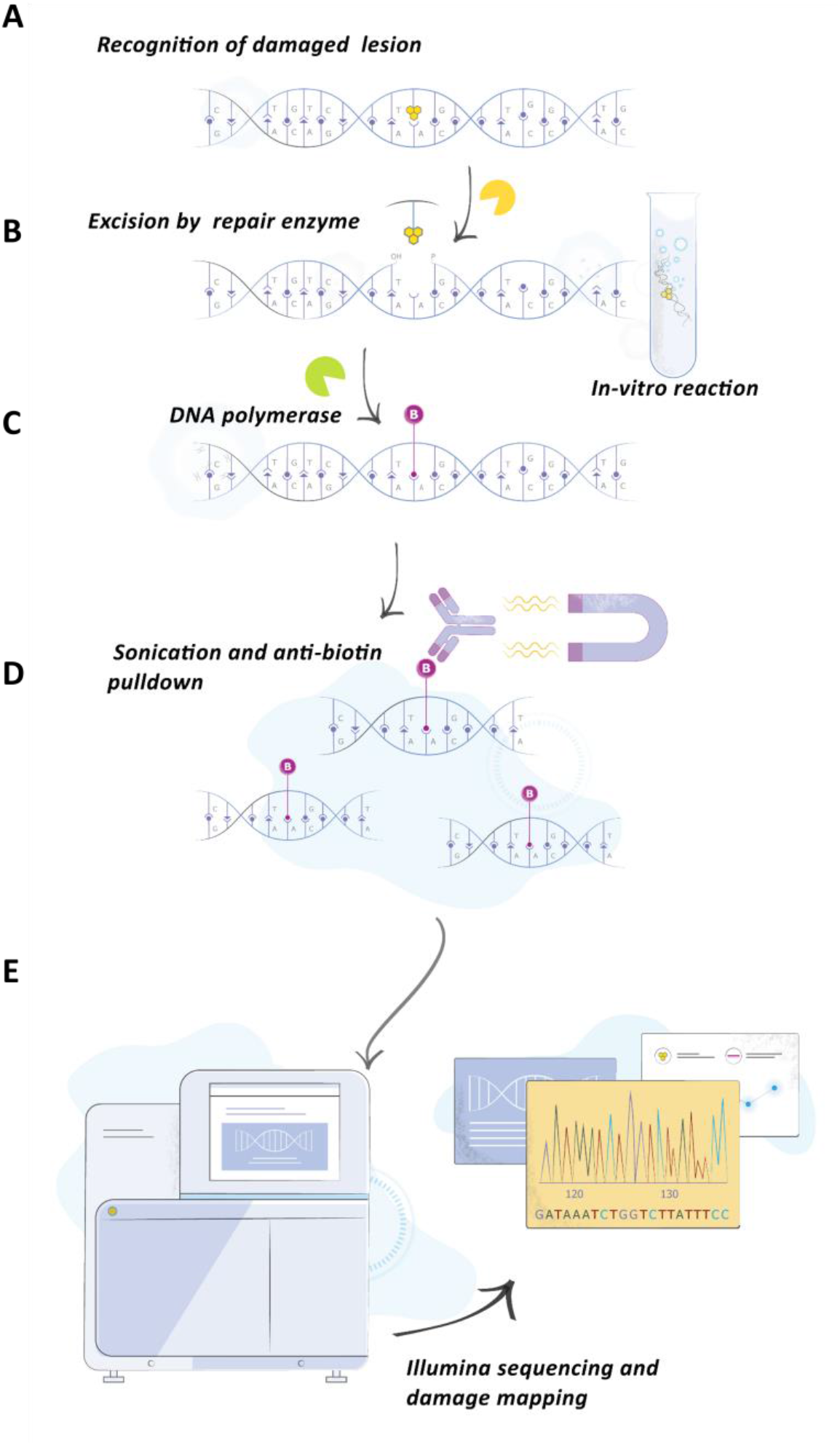
RADD-seq workflow. **(A)** DNA damage lesion is recognized by a specific repair enzyme. **(B)** The damaged lesion is excised, creating a gap in the DNA chain. **(C)** DNA polymerase fills the created gap with biotinylated nucleotides. **(D)** DNA is fragmented to ~150 bp fragments, and pulled down by anti-biotin antibody coated magnetic beads **(E)** Enriched DNA is processed for sequencing and mapped to the reference genome. The damage level across the genome is quantified by assessing the number of mapped reads.

First, validation of RADD-seq was achieved by showing that damaged regions are specifically detected, enriched and accurately sequenced using this method. The sequence motif-specific Nt.BspQI nicking enzyme was used to induce sequence-specific single-strand breaks in genomic DNA extracted from human keratinocyte cells. A map with the expected damage sites was generated from the human genome reference according to the nicking enzyme’s recognition sequence motif (GCTCTTCN^). Nt.BspQI damaged DNA was repaired using biotinylated nucleotides and subsequently pulled down and utilized for library construction and sequencing (**Fig. 2A**). As shown in **Fig. 2B**, the damage sites revealed by RADD-seq correlated well with the expected nicking sites, demonstrating the ability of the method to construct reliable DNA damage maps on a genome-wide scale. With an average peak coverage of 30X, the median of the mapping resolution of single-strand breaks was ~20 bp (**Supplementary Fig. S1**).

**Figure 2.**
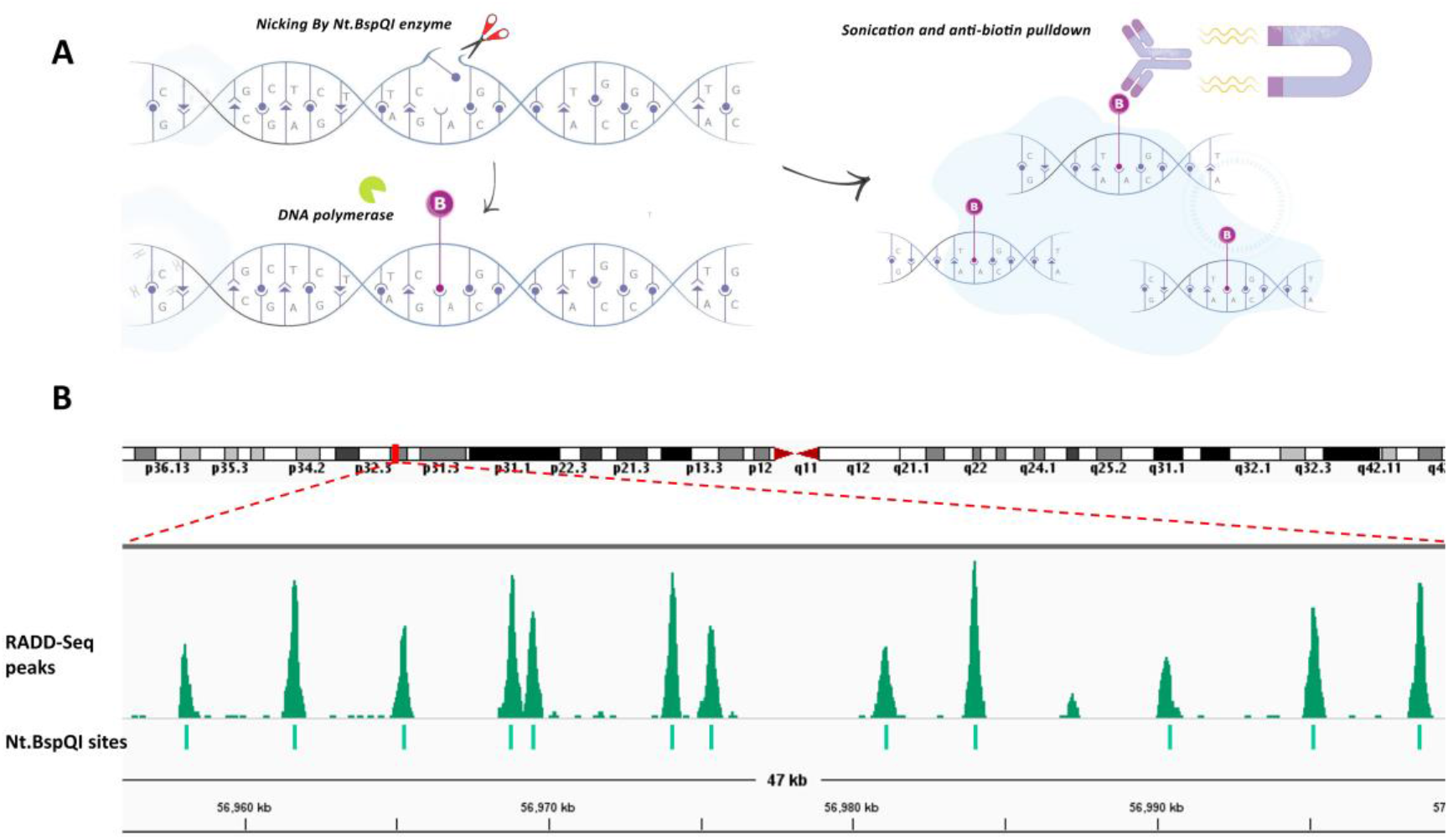
RADD-seq identifies damage sites correctly. **(A**) Illustration of the proof-of-concept experiment. The nicking enzyme Nt.BspQI creates expected nicks along the genome. DNA polymerase is then used to fill the created nicks with biotinylated nucleotides. Finally, DNA is fragmented and pulled down, and the enriched fragments are sequenced. **(B)** Comparison between high coverage regions in the Nt.BspQI sample (top) and the expected nicking site positions of Nt.BspQI (bottom).

### Oxidative damage is heterogeneously distributed

Following its validation, we utilized RADD-seq to reveal DNA damage and repair dynamics in a biological model system. Little is known about the distribution of oxidative DNA damage throughout the human genome. RADD-seq has the potential to link DNA damage to environmental exposures and to specific biological processes such as gene expression and chromosomal packing. To demonstrate this, oxidative DNA damage was induced by KBrO_3_, which generates 8-oxoG in the context of GG and GGG sequences (Kawanishi and Murata 2006). U2OS cells were exposed to 50 mM KBrO_3_ for 1 h. The genomic DNA was extracted, and RADD-seq performed using the oxidative damage repair enzyme hOGG1.

The genome-wide damage distribution was obtained from the number of reads aligned to each genomic locus after normalizing to data from a DNA sample that was not subjected to KBrO_3_ (see Materials and Methods section). The genome-wide oxidative DNA damage distribution is illustrated in **Fig. 3A**. Evidently, oxidative damage hotspots are distributed heterogeneously along the genome indicating that damage induction is not random.

**Figure 3.**
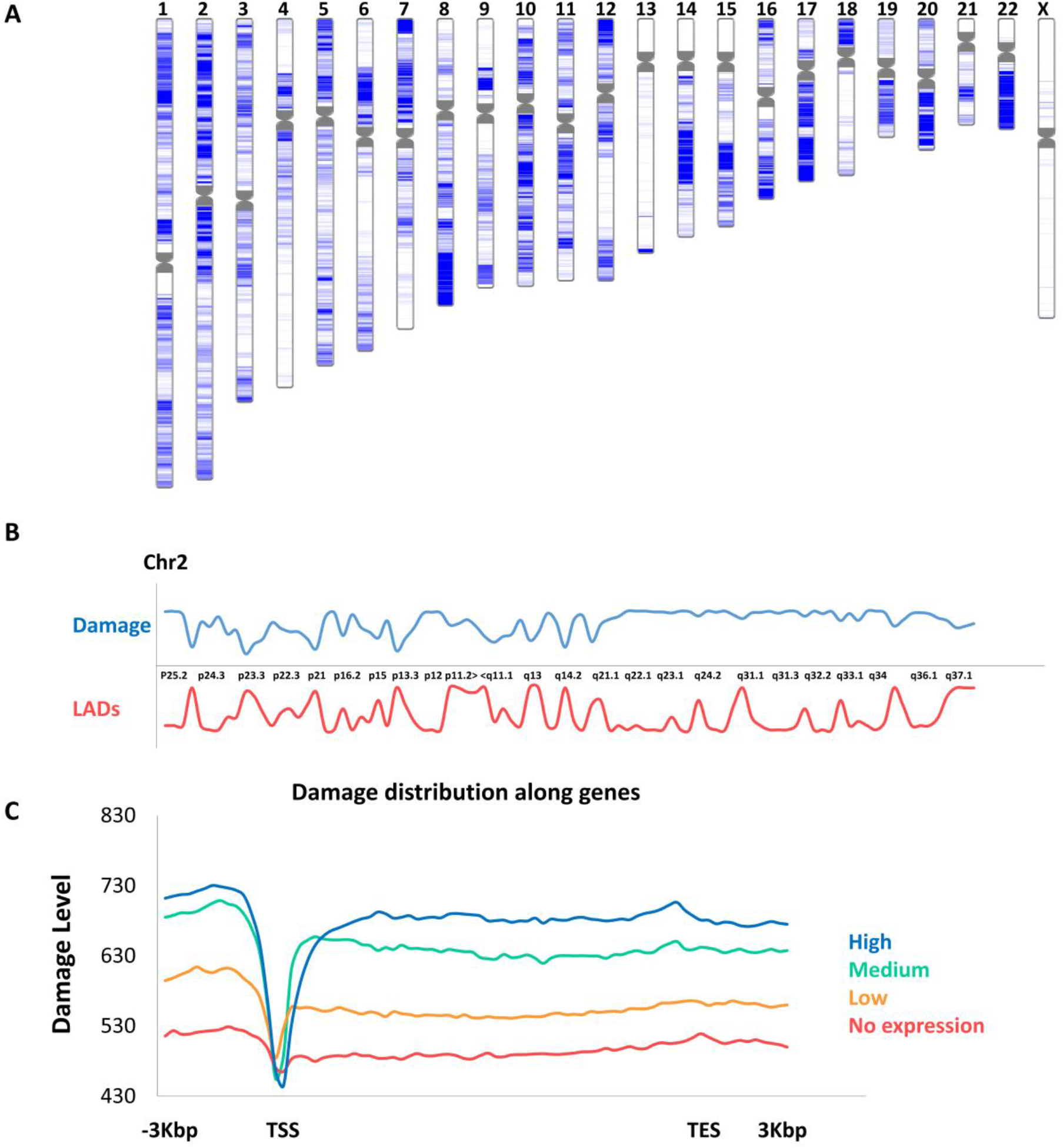
DNA damage distribution revealed by RADD-seq. **(A)** Chromosomal view of the oxidative DNA damage landscape, where darker blue color represents areas with increased damage level. **(B)** A zoomed-in view of the oxidative DNA damage distribution along chromosome 2 (blue) alongside the distribution of lamina-associated domains (LADs, red). **(C)** Oxidative DNA damage in genes with different expression levels, namely high (20% top expressed genes, blue), medium (20% genes with average expression level, green), low (20% bottom expressed genes, yellow), or no expression (red).

Interestingly, when comparing the DNA damage distribution along the genome to the locations of lamina-associated domains (LADs), an anti-correlation of −66% is found. LADs are condensed areas in the chromatin, found in close contact with the nuclear lamina (Friedberg 2003; Van Steensel and Belmont 2017). **Fig. 3B** shows the anti-correlation between the oxidative damage and the LADs distribution along chromosome 2. Hence, oxidative DNA damage is less abundant in these more compacted areas. Moreover, DNA damage accumulation occurred along gene bodies and exhibits a drastic decrease near transcription start sites (TSS, **Fig. 3C**), in line with the findings of Amente and co-workers (Amente et al. 2019). We hypothesize that the decrease in damage at the TSS is due to the tendency of these areas to attract multiple DNA-binding proteins, which may shield the TSS sequence from damaging agents. When inspecting the damage level in genes with different expression levels, we found that higher the expression levels correlate with more damaged gene bodies (**Fig. 3C**).

### Gene expression and chromatin state effect repair dynamics

We further explored the application of RADD-seq for gaining insights into the DNA repair process. U2OS cells exposed to 50 mM KBrO_3_ for 1 h were compared to cells that were washed and allowed to repair for 1 h post damage induction. Genomic DNA was extracted from both groups and subject to RADD-seq (**Fig. 4A**).

**Figure 4.**
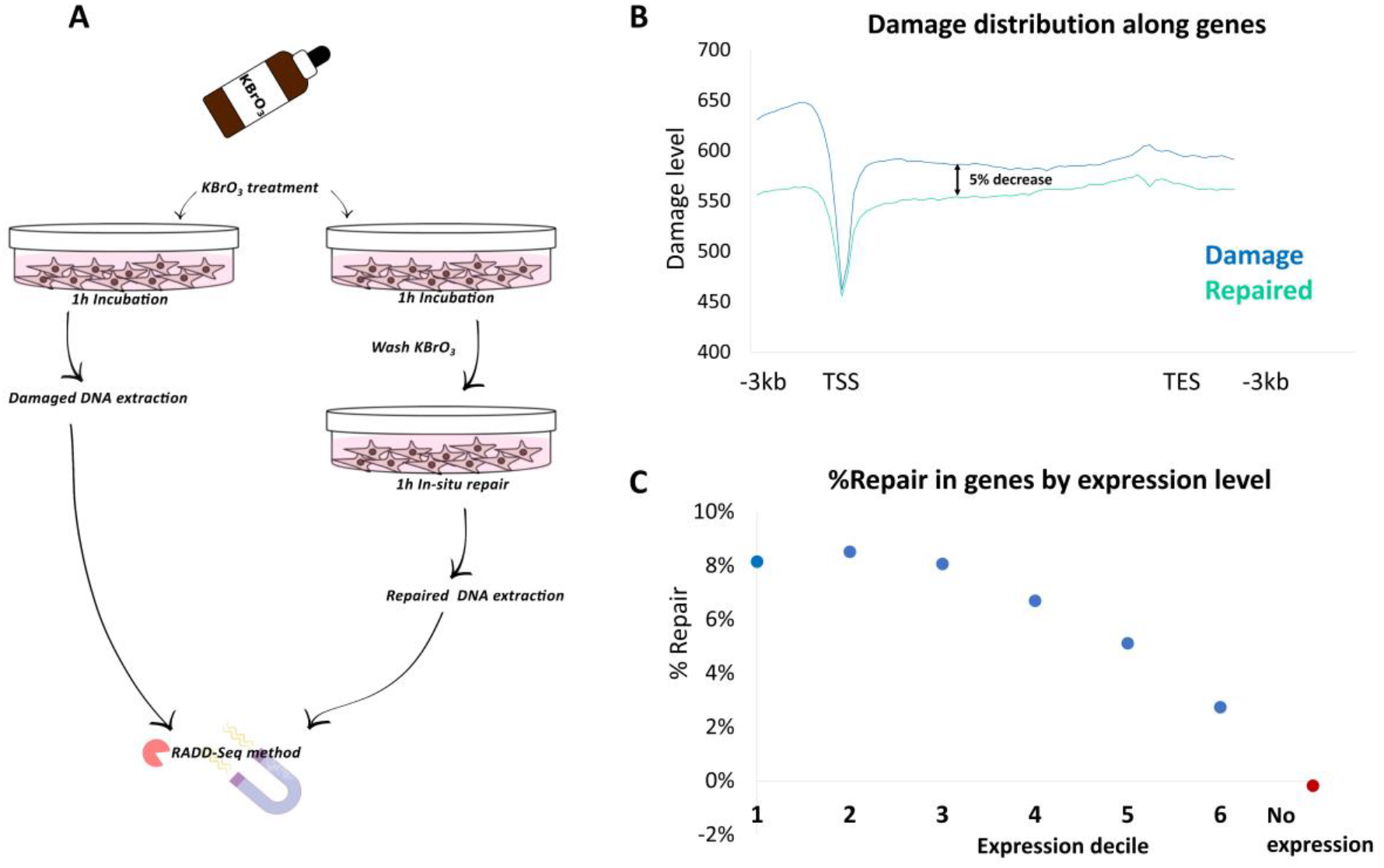
DNA repair of oxidative damage revealed by RADD-seq. **(A)** Illustration of repair dynamics experiment. U2OS cells were treated with 50 mM KBrO_3_ for 1 h, followed by immediate DNA extraction (damage samples), or washed and left for 1 h to repair before DNA extraction (repaired samples). **(B)** Oxidative DNA damage level along all human genes in the damage sample (blue) and the repaired sample (green). **(C)** The average repair level in genes according to their expression levels. Each dot represents the averaged repair level of the specific gene decile (~2,000 genes per dot).

We first evaluated the global DNA damage levels in the two samples using an optical method known as Rapid-RADD (Gilat et al. 2020). The optical experiment showed a ~30% decrease in the global damage level in the repaired samples compared with the damaged samples, verifying that DNA is indeed repaired during the 1h experimental repair period (**Supplemental Fig. S2**). Briefly, both types of samples were treated with a cocktail of repair enzymes specific to oxidative DNA damage, followed by the addition of DNA polymerase that incorporates fluorescent nucleotides into the formed gaps. The labeled DNA was deposited onto a partitioned poly-L-lysine coated glass slide, which was scanned using a slide scanner to evaluate the damage level in each well. In contrast to Rapid-RADD, that quantifies DNA damage in native unamplified DNA, RADD-seq did not show a significant difference in the global number of unique reads mapped for each sample. Given the relatively low sequencing depth used for this experiment and the PCR amplification step during library preparation, cells that were not allowed to go through repair (termed “damage samples” hereafter) differed by only 0.7% in the global number of mapped reads as compared with cells that were allowed to repair (termed “repaired samples” hereafter). Nevertheless, RADD-seq was able to reveal repair patterns in specific regions of the human genome. When comparing gene bodies, we measured an average of 5% decrease in damage level for repaired samples (**Fig. 4B**). More interestingly, when grouping genes by expression level deciles, we found clear positive correlation between repair level and gene expression level, where a higher expression level is associated with a higher level of repair (**Fig. 4C, Supplemental Fig. S3**).

In light of the observed relation between LADs and damage accumulation, we were motivated to observe the repair occurring in these areas. In general, a significant decrease in the damage level was found inside LADs compared with regions up and downstream to LADs for both damaged and repaired samples, (**Fig. 5A**). Moreover, the damage level inside LADs in the damage sample is lower than its corresponding genome-wide average. Interestingly, while outside the LAD regions damage levels decrease after repair, the damage level inside LADs is higher in the repaired sample; presumably, due to the small size of the damaging KBrO_3_ molecules, which may penetrate through condensed chromatin areas such as LADs and accumulate with time. Furthermore, damaged LADs may not be easily accessible for the cellular DNA repair machinery, which is composed of bulky proteins.

**Figure 5.**
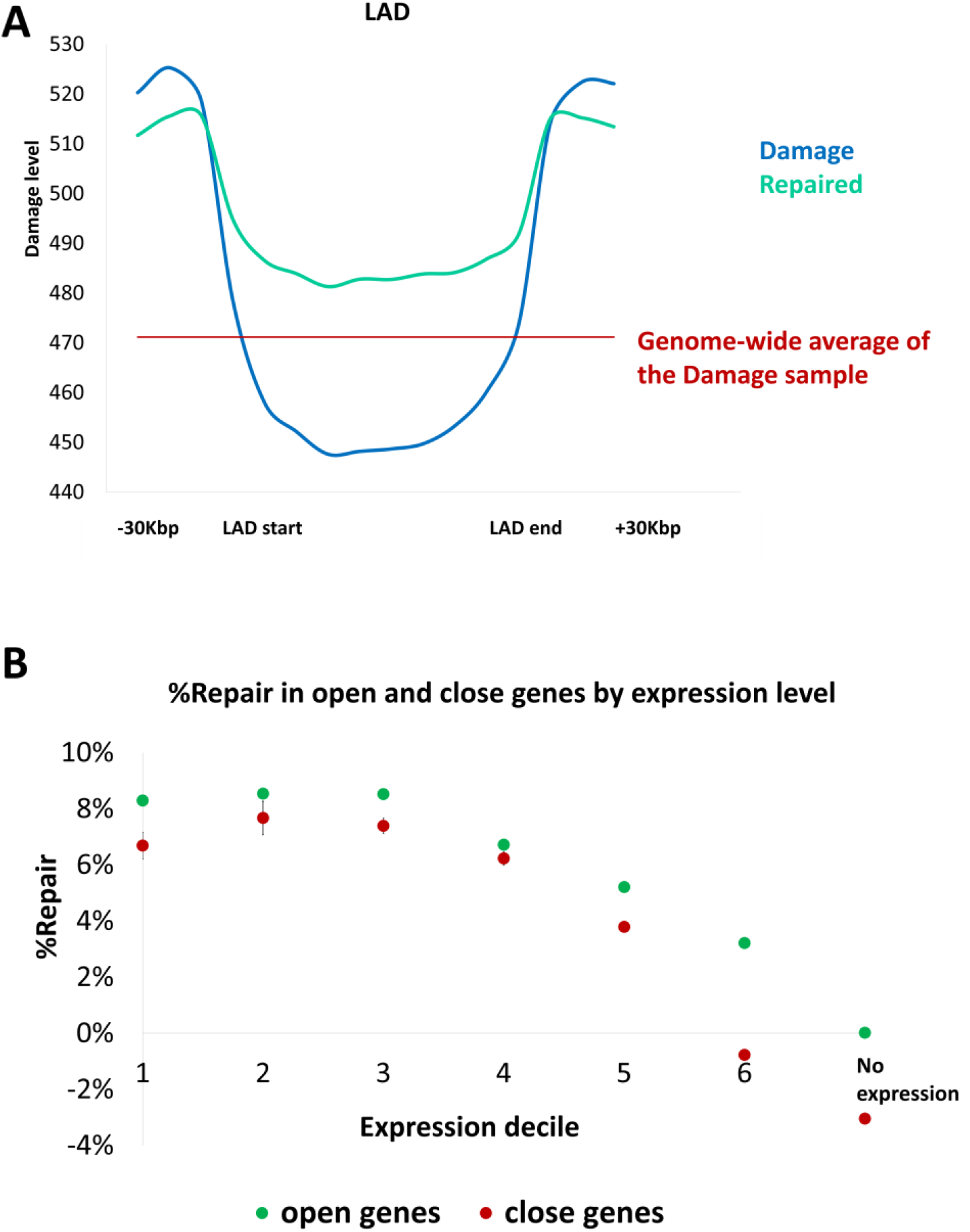
Repair in closed vs. open chromatin genomic regions by RADD-seq analysis. **(A)** The averaged damage level in and outside of LADs in the damage sample (blue) and repaired sample (green). Red line represents the average genome-wide damage level in the damage sample. **(B)** The averaged repair level in genes with different expression levels, divided to open-chromatin genes (green) and closed-chromatin genes (red). Each dot represents the averaged repair level of the specific gene decile.

Next, we examined whether the level of repair in gene bodies is affected not only by gene expression, but also by the state of chromatin. To this end, we divided each of the above-mentioned expression groups into open chromatin or closed chromatin genes. **Fig. 5B** shows the repair level in each group across expression deciles. The repair level in open chromatin genes is higher than in closed chromatin genes from the same decile. Furthermore, in the case of low-expression genes, the difference in repair between the two groups becomes more distinct.

### Essential genes are repaired more extensively

We next evaluated the relation between DNA repair and functionally different gene groups (**Fig 6A**). Generally, genes essential for basic cellular functions tended to be more extensively repaired than other genes. Housekeeping and translation-related genes showed the highest repair levels. **Fig. 6B** shows an example of the damage level before and after repair for a housekeeping gene, CDK1 (top), and a translation-related gene, EIF4A2 (bottom). These data clearly demonstrate the distinct reduction in damage level in these genes following repair. In contrast, genes related to the nervous system and smell perception, not essential for the function of U2OS cells, presented a reduced level of repair (**Fig 6A**).

**Figure 6.**
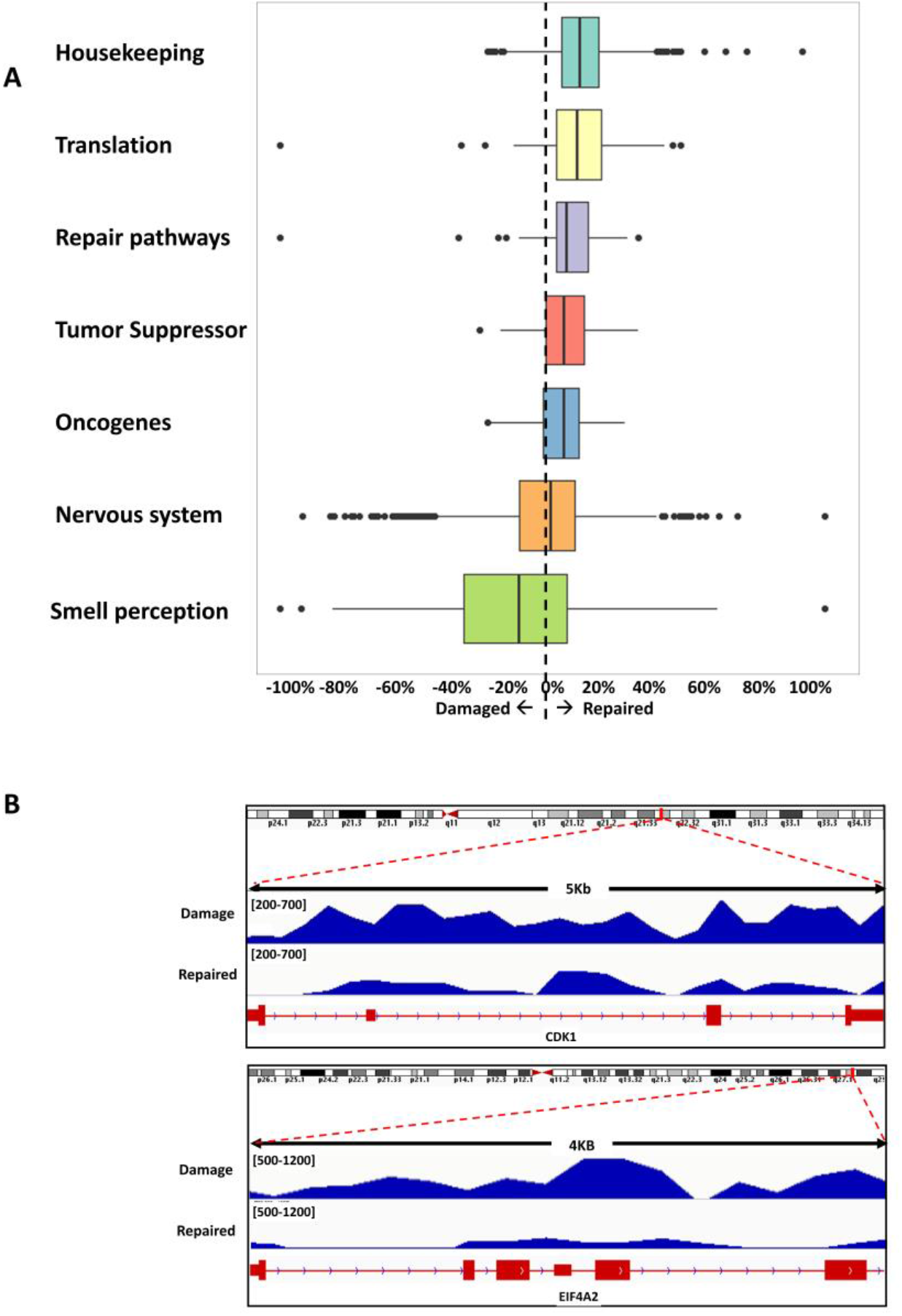
Repair in Gene groups. **(A)** Box plot comparison of the repair level in six different gene groups; housekeeping genes (mean repair: 12.5%, number of genes: 931), translation related genes (mean repair: 12.5%, number of genes: 160), repair pathway genes (mean repair: 8.8%, number of genes: 57), tumor suppressor genes (mean repair: 6.8%, number of genes: 63), oncogenes (mean repair: 6.4%, number of genes: 82), nervous system genes (mean repair: −3.4%, number of genes: 1286) and smell perception genes (mean repair: −11.3%, number of genes: 384). **(B)** Representative damage and repaired signals of a housekeeping gene (CDK1, top) and a translation-related gene (EIF4A2, bottom) as obtained by RADD-seq analysis.

We note that most of the genes in the top-repaired groups, housekeeping genes and translation-related genes, belong to the top gene expression decile. To evaluate the impact of gene expression level on DNA repair level, we compared these two most highly repaired gene groups with the rest of the genes in the top decile of gene expression level. We found that the repair level of genes belonging to these gene groups was significantly higher than the overall repair level of the rest of the genes with the same gene expression level, indicating that additional factors influence the level of repair (**Supplemental Fig. S4**).

Finally, we show that the highest and lowest repaired gene groups include genes with specific functionalities. The top 500-repaired genes consist of a large group of translation-related genes and a group of cell-cycle related genes (both part of the housekeeping gene group, **Supplemental Fig. S5A**). The bottom 500-repaired genes (which mostly include genes with a higher post-repair damage level) consist of two large clusters of genes, one involving thiol-dependent ubiquitinyl hydrolase activity, and another, related to smell perception (**Supplemental Fig. S5B**). These findings are in line with the fact that the studied cells originate from bone tissue, which unlikely requires the activity of the bottom 500-repaired genes.

## Discussion

Oxidative DNA damage poses a serious threat to genome integrity and has been linked to the development of cancer, neurodegeneration and cell senescence (Cadet and Davies 2017; Whitaker et al. 2017; Barnes et al. 2019; Maiuri et al. 2019). Yet, to date, little is known about the genome-wide distribution of oxidative DNA damage loci and the repair dynamics of this damage type. RADD-seq provides a rapid and inexpensive mapping of DNA damage and repair based on pull-down and sequencing technique. Oxidative DNA damage map was obtained with as little as 60 million reads per sample. Damage was found to accumulate in various regions along the human genome, and was more abundant in highly expressed genes and in regions with less condensed chromatin. We found that repair tends to occur primarily for highly expressed genes. Moreover, specific gene groups, vital to the normal functioning of the cells such as housekeeping genes, were repaired more considerably than other, less crucial genes. This last notion indicates that not only physical properties, such as chromatin architecture, direct DNA repair enzymes to their repair targets, but also other factors, yet unknown. Both basic research and clinical practice rely upon understanding the type and extent of genomic DNA damage. The detection of damage hotspots can help determine disease predisposition, whereas the targeting of areas with high repair levels can aid in monitoring therapeutic response. From an analytical perspective, RADD-seq offers an adaptor-free amplification for a variety of DNA damage types. Coupled with Rapid-RADD for global quantification, these two methods provide means for determining both quantities and locations of DNA damage, the combination of which is essential for determining dose-response relationships of DNA-damaging agents. Here, we demonstrate RADD-seq applicability toward oxidative DNA damage; however, by choosing different repair enzymes, this protocol can be easily adjusted for other types of DNA lesions.

## Methods

### Cell culture

U2OS (human osteosarcoma) cells were cultured in Dulbecco’s modified Eagle’s medium (DMEM), supplemented with 10% fetal bovine serum (Gibco), L-glutamine (2 mM) and 1% Penicillin-Streptomycin (10,000 U/mL, Gibco). Cells were incubated at 37 °C with 5% CO_2_.

### KBrO_3_ treatment

Final concentration of 50 mM KBrO_3_ was added to culture medium for one hour. Cells were washed with PBS twice, and genomic DNA was immediately extracted (for damage samples). Alternatively, cell medium was replaced, and cells were allowed to repair for one hour at 37 °C with 5% CO_2_ before DNA extraction (for repaired samples). Cells were allowed to grow up to a week post KBrO_3_ treatment, to validate their viability.

### DNA extraction

DNA was extracted from approximately 10^6^ cells using the “GenElute-Mammalian Genomic DNA Miniprep Kit” (Sigma) according to manufacturer’s instructions.

### Labeling oxidative DNA damage

KBrO_3_-treated DNA samples were labeled for oxidative damage in three consecutive enzymatic reactions. In the first step, each reaction tube contained 2.2 μg of DNA sample, 2 μL of 10x buffer 4 (New England Biolabs, NEB), 2 μL of 1 mg/ml bovine serum albumin solution (NEB), 1 μL of hOGG1 (ProSpec TechnoGene Ltd.) and ultrapure water to a final volume of 20 μL. The reaction mixture was incubated for 30 minutes at 37 °C. In the second step, 1 μL of Endonuclease IV (10,000 U/ml, NEB) was added to the reaction, and it was incubated for additional 30 minutes at 37 °C. In the final step, the following were added into each reaction tube: 0.5 μL of 10x buffer 4 (NEB), 0.5 μL of 1 mg/mLbovine serum albumin solution (NEB), 1.2 μL of 50 mM NAD^+^ (NEB), deoxynucleotides (A,G,C (sigma) and biotinylated-dUTP (ThermoFisher Scientific)) to a final concentration of 100 nM, 0.5 μL of Bst DNA polymerase, Large fragment (8,000 U/mL, NEB), 0.5 μL of Taq DNA polymerase (5,000 U/mL, NEB), 0.4 μL of Taq DNA ligase (40,000 U/mL, NEB) and ultrapure water to a final volume of 30 μl. The reaction mixture was incubated for 30 minutes at 65 °C.

### DNA fragmentation

Following damage labeling, 120 μL hydrated DNA was transferred to a Covaris microtube (AFA fiber pre-slit snap cap, 6×16 mm) and sheared using a probe sonicator (Covaris S220 focused ultrasonicator instrument, operation system SonoLab 7.0, pre-chilled to 4 °C, and degassed for 30 minutes). Samples were sonicated for 10 minutes at a peak power of 175 W, duty factor 20%, and 200 cycles per burst. The target fragmentation size was 150 bp. The fragmented and labeled DNA samples were purified from excess nucleotides using “QIAquick PCR Purification Kit” columns (QIAGEN), according to manufacturer’s recommendations.

### Immunoprecipitation

Fragmented DNA was immunoprecipitated using anti-biotin antibodies (1 mg/mL, abcam) and protein G beads (Invitrogen). 70 μL of protein G beads were washed twice with 70 μL IP buffer (10 mM Tris pH 7, 1 mM EDTA, 150 mM NaCl), and were resuspended in 70 μL IP buffer. The 70 μL protein G beads were divided into two tubes: (1) 50 μL beads were incubated with 5 μg of anti-biotin antibodies (2) 20 μL beads were incubated with 500 μL of the fragmented DNA (to allow nonspecific DNA binding to the beads). Both tubes were incubated for 2 h at 4 °C with rotation and vibration. Following incubation and magnetic pull-down for both tubes, the supernatant from tube 1 was discarded, and the supernatant from tube 2 was added to the beads in tube 1. The beads and DNA were incubated overnight at 4 °C with rotation and vibration. Following incubation, the beads were washed seven times with 700 μL of IP buffer for five minutes at 4 °C with rotation and vibration. To elute DNA, the beads solution was incubated with 40 μL elution buffer (10 mM TE buffer, 1 mg/mL proteinase K and 0.5% SDS), for three h at 50 °C with shaking at 900 rpm, followed by magnetic pull-down. The pulled-down DNA was purified using “QIAquick PCR Purification Kit” columns (QIAGEN), according to manufacturer’s recommendations.

### Proof of concept experiment with Nt.BspQI nicking enzyme

DNA from human keratinocyte cells was extracted using the “GenElute-Mammalian Genomic DNA Miniprep Kit” (Sigma) according to manufacturer’s instructions. DNA was nicked using the Nt.BspQI enzyme (NEB, 10,000 U/mL). 10 μg of DNA were mixed with 11.6 μL Nt.BspQI enzyme, 10 μL buffer 3.1 (NEB) and ultrapure water to a final volume of 70 μL. The reaction mixture was incubated for 2 h at 50 °C. Following nicking reaction, DNA samples were labeled with 10 μL of a cocktail of repair enzymes (PreCR mix, NEB), fragmented, and assayed according to RADD-seq procedure as described above.

### Next generation sequencing

Sequencing libraries were prepared using TruSeq DNA Nano library preparation kit (Illumina) according to manufacturer’s instructions, without additional fragmentation and without size-selection. All sequencing libraries were sequenced on HiSeq 2500 platform by the Technion Genome Center. Two biological replicates of each data type; damage and repaired, were prepared for sequencing, as well as two replicates with no repair enzyme as controls for each reaction. In addition, two replicates of input DNA, containing non-damaged DNA were prepared. For each data type, 62M 50 bp single reads were collected.

### Sequence analysis

Sequencing reads were aligned to the hg38 human reference using Bowtie 2 (version 2.3.4.2) with default parameters. Following alignment, reads with MAPQ less than 30 were filtered with SAMtools, and the remaining reads were de-duplicated with MateCigar (version 2.18.2.1) to eliminate PCR duplication bias. Genomic coverage damaged DNA was calculated using BEDTools genomecov (version 2.27.1). All software mentioned above were used through the “UseGalaxy” site (https://usegalaxy.org/). Each of the data sets, damage, repaired and input data, were generated using an in house sliding window script with 200 bp window size and 1bp length step size. Correlation of biological replicates was assessed using Pearson correlation in 1 kb bins (**Supplemental Fig. S6**). The two biological replicates for damage and repair data sets were combined by averaging the number of reads in each region. The data in both sample types (“damage” and “repaired”) was normalized to the input DNA, to eliminate any bias resulting from the pull-down assay and mis-alignment to repetitive areas in the genome. In the first step, we divided the input DNA into 200 bp sized-windows, and the read count in each window falling in the range of (mean+1Stdv - mean+2Stdv) was divided by mean+1Stdv (values were ranging between 1-1.14). In the second step, values lower than mean+1Stdv was set to 1, and all regions higher than mean+2Stdv were discarded from the data since they represent areas with high background level. Next, the read count for damage and repaired data in each 200 bp window was divided by the normalized input DNA respective windows. The regions discarded from the input data were discarded from these data sets as well. Finally, reads mapping to the ENCODE-defined blacklist regions (https://sites.google.com/site/anshulkundaje/projects/blacklists) were discarded. This normalization method normalizes only regions that aligned to regions with read count of mean+1Stdv – mean+2Stdv in the input DNA. All the regions below the mean value fall into the background noise level, and all the areas above the mean+2Std are considered outliers and mainly represent repetitive areas in the genome.

### Correlation of DNA damage and repair to gene expression and chromatin state

Gene expression was assessed using RNA-seq data of U2OS cells, obtained from NCBI BioProject (accession number PRJNA668283). For the damage analysis, genes were classified into four groups: high expression (20% top expressed, ~4,000 genes), medium expression (~4,000 genes with average expression level), low expression (20% bottom expressed, ~4,000 genes), and no expression (FPKM=0, ~3,000 genes). For the repaired analysis, genes were classified into six expression deciles (each with ~2,000 genes) and another group of no expression gene (FPKM=0, ~3,000 genes).

Open and closed chromatin regions were determined using ATAC-seq data of U2OS cells, obtained from NCBI BioProject (accession number PRJNA486188). Genes were divided into two groups based on their “openness” level: genes in which over 50% of the gene body was found in open chromatin areas were defined open (~16,000 genes), the rest of the genes were defined as closed (~4,000 genes).

### Gene ontology analysis

Gene ontology “GO” analysis was performed with the web-based tools Panther Gene Ontology Consortium’s web tool (http://pantherdb.org/) and STRING (https://string-db.org/). Sorting genes into different gene groups was done using online databases (Housekeeping-https://www.tau.ac.il/~elieis/HKG/, repair pathways-https://www.mdanderson.org/documents/Labs/Wood-Laboratory/human-dna-repair-genes.html, oncogenes and tumor suppressor-https://cancerres.aacrjournals.org/content/canres/suppl/2012/01/23/0008-5472.CAN-11-2266.DC1/T3_74K.pdf, translation-related, smell perception, nervous system – Panther Gene Ontology).

### Optical validation experiment (Rapid-RADD)

DNA samples extracted for the RADD-seq protocol, were used for Rapid-RADD procedure as well as described by Gilat et al (Gilat et al. 2020). In short, KBrO_3_-treated DNA samples were labeled for oxidative DNA damage as described above, where instead of adding biotinylated-dUTP, a fluorescent ATTO-550-UTP (Jena biosciences GMBH) in the same quantity was added in the nucleotide mixture. Next, labeled DNA samples were purified from excess fluorophores using “Oligo Clean & Concentrator” columns (Zymo research), according to manufacturer’s recommendations, with two washing steps for optimal results. Upon preparing the multi-well slide according to Rapid-RADD protocol, 1 μL of labeled-DNA samples were placed in each well, and incubated according to protocol instructions. The slide was scanned, stained for total-DNA quantification, and scanned again, and results were analyzed to quantify the amount of damage in each sample.

### Data Access

All raw and processed sequencing data generated in this study have been submitted to the NCBI SRA BioProject & Biosample (SRA; https://www.ncbi.nlm.nih.gov/sra/) under accession number PRJNA697255.

## Acknowledgments

This study was funded by the European Research Council Consolidator grant (Grant No. 817811, YE) and the US National Institute of Health (Grant R21ES028015, NRG and YE).

## Author Contributions

Y.E. and Y.M conceived and supervised the project. N.G, Y.M and H.S performed the experiments. All authors analyzed the data. All authors wrote and edited the manuscript. N.G and D.F contributed equally to this work.

