## Supplementary for "Genome wide mapping of DNA lesions by Repair Assisted Damage Detection sequencing – RADD-Seq"

**Contents**

**Supplemental Figures**

**Supplemental Figure S1** …..…………………………….……………………………………... 2

**Supplemental Figure S2** …..…………………………..………………………………………... 2

**Supplemental Figure S3** …..…………………………..………………………………………... 3

**Supplemental Figure S4** …..…………………………..……………………..………………… 4

**Supplemental Figure S5** …..……………………………………………………….………….... 5

**Supplemental Figure S6** …..………………………………………………..…………………….. 6

**References** ……………..…..…………………………………………..…………………… 6

**Figure S1**

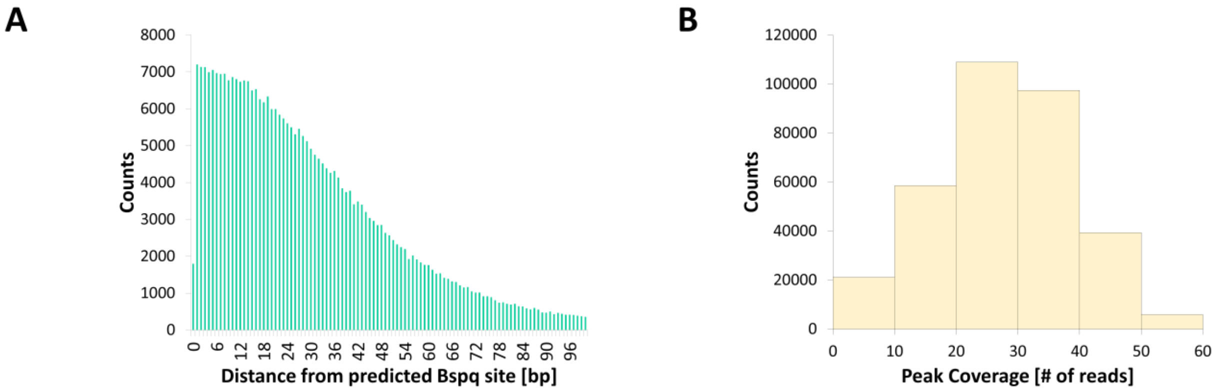


**Figure S1. A proof of concept experiment**. **(A)** Histogram of the distance of high coverage peaks in the Nt.Bspq sample, from the closest Nt.Bspq site.**(B)** Histogram of the peak coverage in the Nt.Bspq experiment.

**Figure S2**

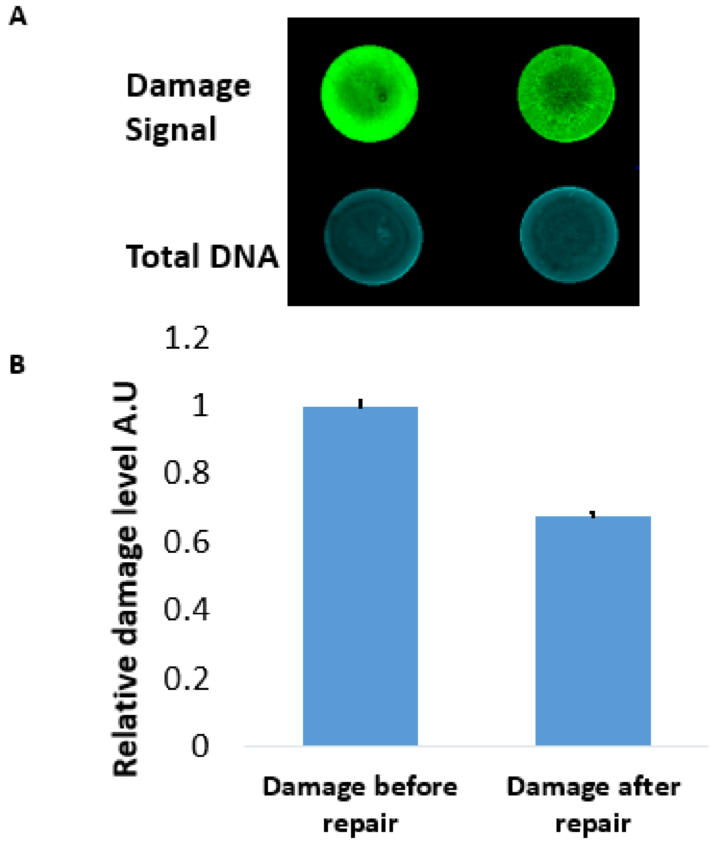


**Figure S2. Rapid-RADD before and after repair of oxidative damage. (A)** U2OS cells were incubated with 50 mM $KBrO_3$, then washed and allowed to undergo native DNA repair. The DNA was extracted immediately post-treatment and 1 hour post-treatment, labeled using the hOGG1 repair cocktail, and further assayed according to the Rapid-RADD protocol (Gilat et al. 2020). Representative fluorescence image of wells for each time point (immediately post-treatment and 1 hour post-treatment); **(B)** Quantification results derived from the slide images and showing significant repair of the damaged DNA. Data represent mean ± standard deviation (n = 5). Each bar represents the averaged results of two independent experiments.

**Figure S3**

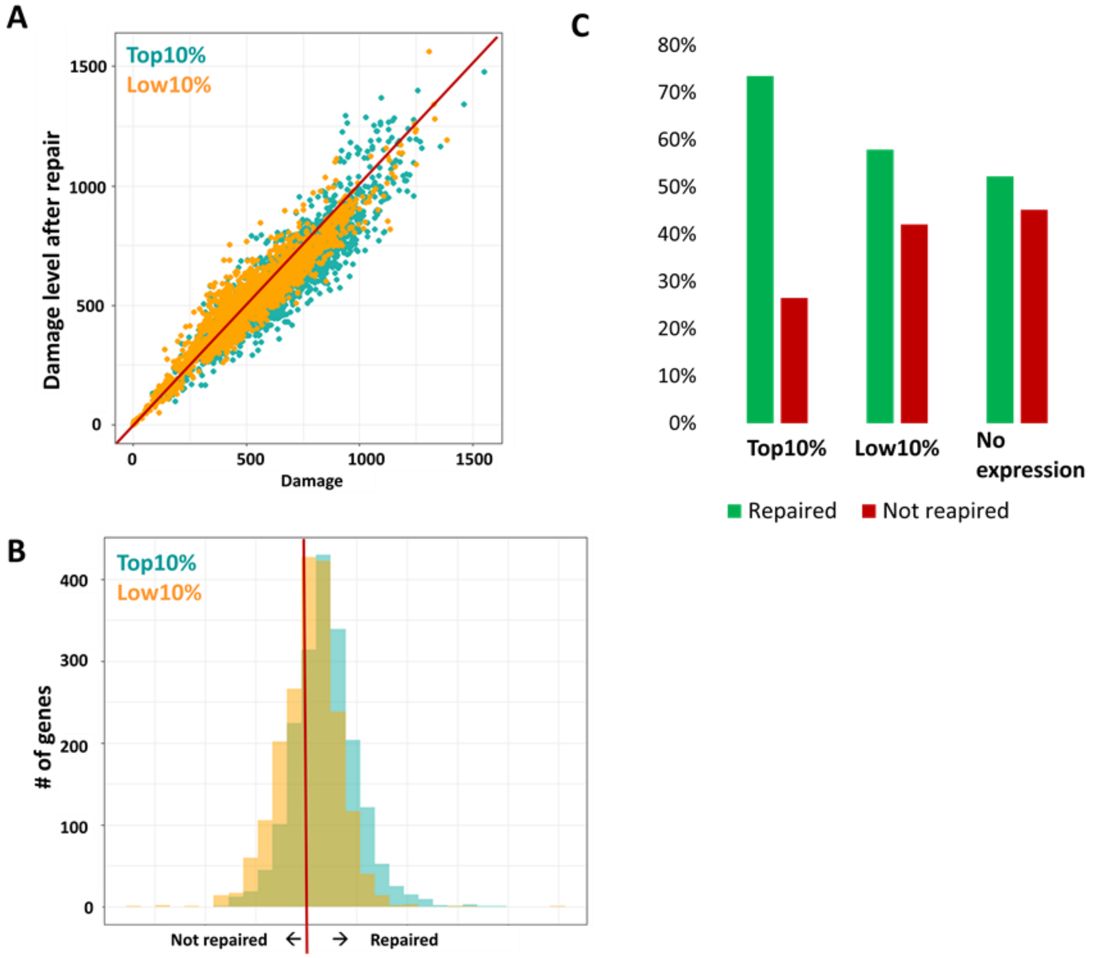


**Figure S3. Visual comparison of repair level in different expression level genes. (A)** Scatter plot of the top 10% expressed genes (turquoise) and the bottom 10% expressed genes (Yellow). **(B)** Histogram of the top 10% expressed genes (turquoise) and the bottom 10% expressed genes (Yellow). **(C)** Bar graph of the top and bottom 10% expressed genes, as well as no expression genes, divided into repaired genes (green) and not repaired genes (red).

**Figure S4**

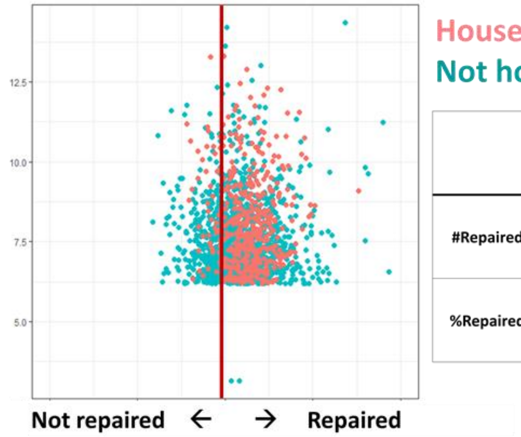


Housekeeping genes
Not housekeeping genes

| | Housekeeping in top 10% expressed | Top10% expressed not including Housekeeping genes |
|---|---|---|
| #Repaired genes | 420 | 1022 |
| %Repaired genes | 86% | 71% |

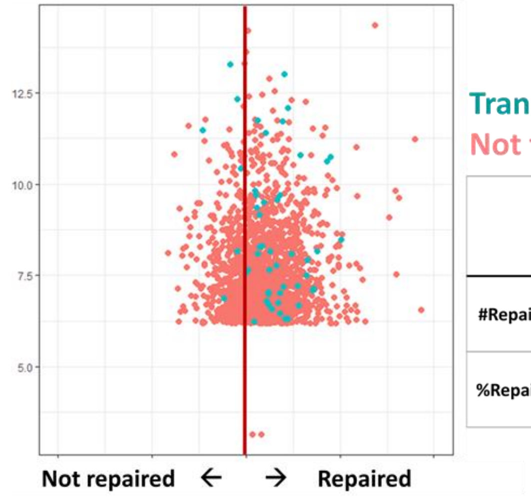


Translation-related genes
Not translation-related genes

| | Translation-related genes in top 10% expressed | Top10% expressed not including translation related-genes |
|---|---|---|
| #Repaired genes | 44 | 1398 |
| %Repaired genes | 88% | 74% |

**Figure S4.** Specific gene groups within the top 10% expressed genes. **(A)** 495 Housekeeping genes are found in the top 10% expressed genes, 86% of these genes are repaired, while only 71% of the top 10% expressed genes not including housekeeping genes are repaired. **(B)** 160 Translation-related genes are found in the top 10% expressed genes, 88% of these genes are repaired, while only 74% of the top 10% expressed genes not including translation-related genes are repaired.

**Figure S5**

**A** **Network enrichment map for the top 500 repaired genes**

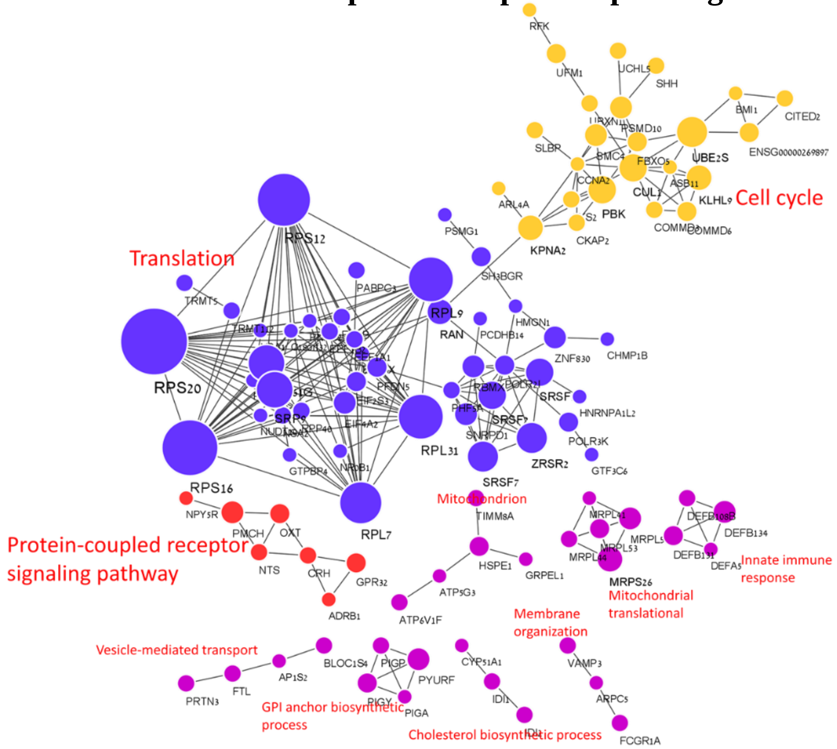


**B** **Network enrichment map for the bottom 500 repaired genes**

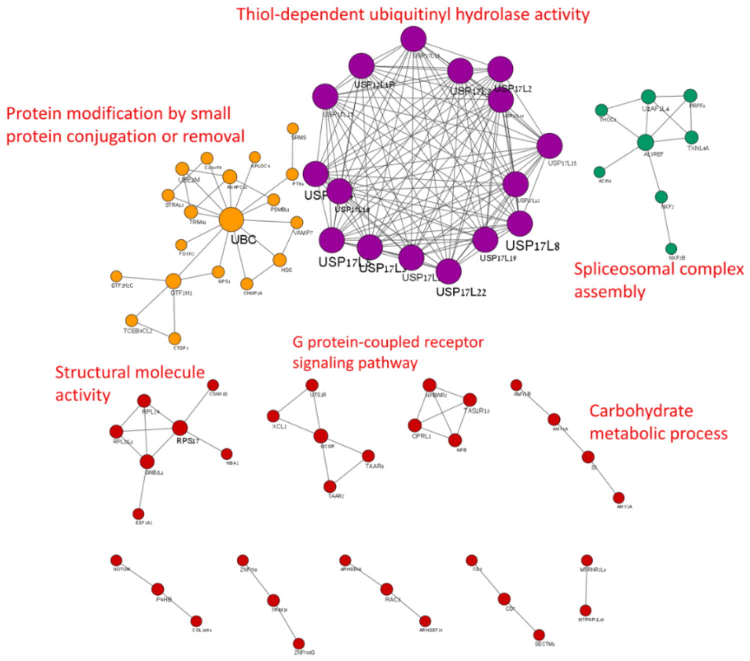


**Figure S5. A Functional Network analysis of genes with different repair levels**. Network enrichment maps analyzed with STRING (https://string-db.org/), showing the relationship between genes which were in the group of top (A) or bottom (B) 500 genes. Genes are represented as colored circles or nodes, and the relationship between them are represented by black lines, or edges. **(A)** A network enrichment map of the top 500 repaired genes. The enrichment p-value in the network is 1.00e-16. **(B)** A network enrichment map of the bottom 500 repaired genes. The enrichment p-value in the network is 2.85e-6.

**Figure S6**

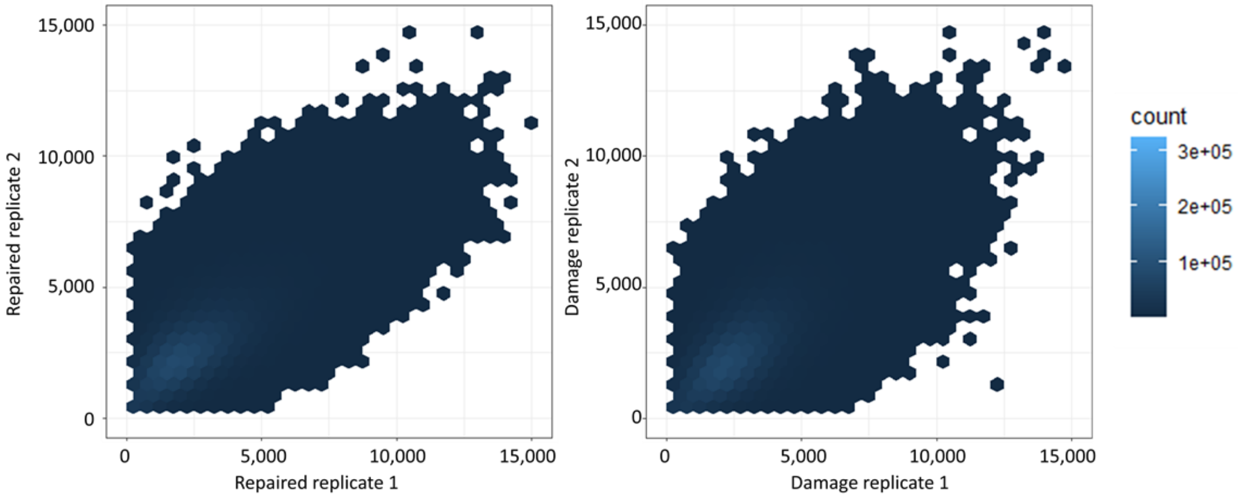


**Figure S6.** The correlation of two biological replicates of the Damage samples (right) and Repaired samples (left) were tested with Pearson correlation test. The correlation was done in 1Kb bins, and was found to be 75% correlation for the Damage replicates, and 76% for the Repaired replicates.
